# Optimized upstream analytical workflow for single-nucleus transcriptomics in main metabolic tissues

**DOI:** 10.1101/2025.02.22.639604

**Authors:** Pengwei Dong, Shitong Ding, Guanlin Wang

## Abstract

Single-nucleus RNA sequencing (snRNA-seq) has emerged as a powerful approach for studying cellular heterogeneity in metabolic tissues. However, snRNA-seq analysis remains challenging due to low gene expression and data complexity. Here, we introduce an optimized analytical workflow for snRNA-seq data from 67 samples across white adipose tissue, muscle, liver and the hypothalamus. We emphasized the importance of key steps including ambient RNA removal, doublet identification and data integration to ensure accurate downstream analysis. This workflow offers a valuable resource for researchers in metabolism, facilitating deeper insights into cellular diversity and metabolic function through rigorous snRNA-seq analysis.

---

Metabolic homeostasis is regulated by a network of organs and tissues, primarily involving adipose tissue, muscle, liver and the hypothalamus, which act as central metabolic regulators. Cellular dysregulation within these tissues substantially associates with metabolic disorders, including obesity, type 2 diabetes, and non-alcoholic fatty liver disease (NAFLD) [1]. Understanding the molecular mechanisms governing metabolic control requires dedicated analysis of physiological and pathological cellular heterogeneity within these tissues. However, single cell level investigations to decipher the complexities of cellular mechanisms remain challenging due to the fragile nature of certain cell types and technical noise within these metabolically active tissues, resulting in limited studies compared to well-characterized atlases in immune cell populations [2].

Single-nucleus RNA sequencing (snRNA-seq) has emerged as an alternative approach to address these challenges [3]. By isolating nuclei rather than whole cells, snRNA-seq eliminates the necessity for extensive tissue dissociation, which is particularly advantageous for vulnerable cells or frozen samples. This approach enables high-resolution transcriptional profiling of metabolically active tissues and frozen tumour samples, revealing insights into cellular diversity and regulatory networks crucial for metabolic function. Despite these advantages, analyzing snRNA-seq data from metabolic tissues still presents unique difficulties. Adipose tissue is composed of large, fragile, lipid-rich adipocytes, along with diverse cell types including preadipocytes, immune cells, and endothelial cells, all integral to energy storage and endocrine functions [4-6]. Similarly, muscle tissue comprises muscle fibres, satellite cells, and interstitial cells, which work together to regulate muscle metabolism [7], whereas liver tissue consists of hepatocytes, Kupffer cells, endothelial cells, and hepatic stellate cells, each with distinct gene expression profiles [8]. The heterogeneous cell populations complicate the transcriptomic analysis due to inherently lower mRNA content in nuclei compared to whole cells. Given these challenges, careful handling and analytical methods are essential for the interpretation of snRNA datasets in metabolic tissues. To overcome the low abundance of transcripts in snRNA-seq datasets, the Bayesian-based normalization method *sctransform* has been developed to improve the identification of key cell types [9]. Another notable difficulty in scRNA-seq and snRNA-seq data analysis is the need for effective data integration and batch correction, especially when combining datasets from multiple samples, time points, or different labs for large consortia studies. Biological variations within and between samples can be masked by technical differences or batch effects. To address this challenge, several methods have been developed including Seurat Canonical Correlation Analysis (CCA), Harmony, reciprocal PCA (rPCA), single cell Variational Inference (scVI), and single cell Annotation Variational Inference (scANVI) etc [10]. Each of these methods has unique strengths and limitations, making it essential to evaluate their performance on a case-by-case basis depending on the complexity and heterogeneity of the data. Thus, addressing these complexities of metabolically active tissues requires a systematic tailored upstream analytical workflow to extract meaningful biological insights from snRNA-seq datasets.

We introduce an optimized analytical workflow designed for snRNA-seq data from multiple metabolic tissues, including white adipose tissue, muscle, liver and the hypothalamus, sourced from both human and murine models. It covers processing steps include count matrix generation, ambient RNA removal, doublet identification, quality control of expression matrix, normalization, data integration and benchmarking, emphasizing the importance of ambient RNA and doublets removal for accurate cell type annotation. We have applied it to analyze 67 samples in 5 datasets (201,411 cells in total, **Supplementary Fig. S1a, Supplementary Information**) using 10x Genomics technologies (**Supplementary Table S1**) to show the performance.

The optimized and comprehensive upstream analyses workflow starts from raw FASTQ files (**Fig. 1a, Supplementary Information**). We used standard Cell Ranger pipeline to generate an expression matrix, followed by applying CellBender [11] to remove ambient RNA and scDblFinder for doublet removal tools [12]. The filtered matrix was then processed with Seurat for downstream analyses including expression matrix quality control (QC) by minimum expressed genes, minimum expression UMIs and mitochondria percentage, normalization, dimensional reduction, clustering and annotation. After these preprocessing steps, 141,113 cells past all the QC steps and used in the downstream analysis (**Supplementary Table S1, Supplementary Information**). We incorporated multiple integration strategies, including Seurat CCA, Harmony, rPCA, scVI and scANVI, followed by systematic benchmarking using scIB [13] to help the users to decide the most appropriate integration method. This workflow ensures the reservation of high-quality data for subsequent cell-type classification and functional analysis, enhancing the interpretability and reliability of findings across heterogeneous metabolic tissues.

**Figure 1.**
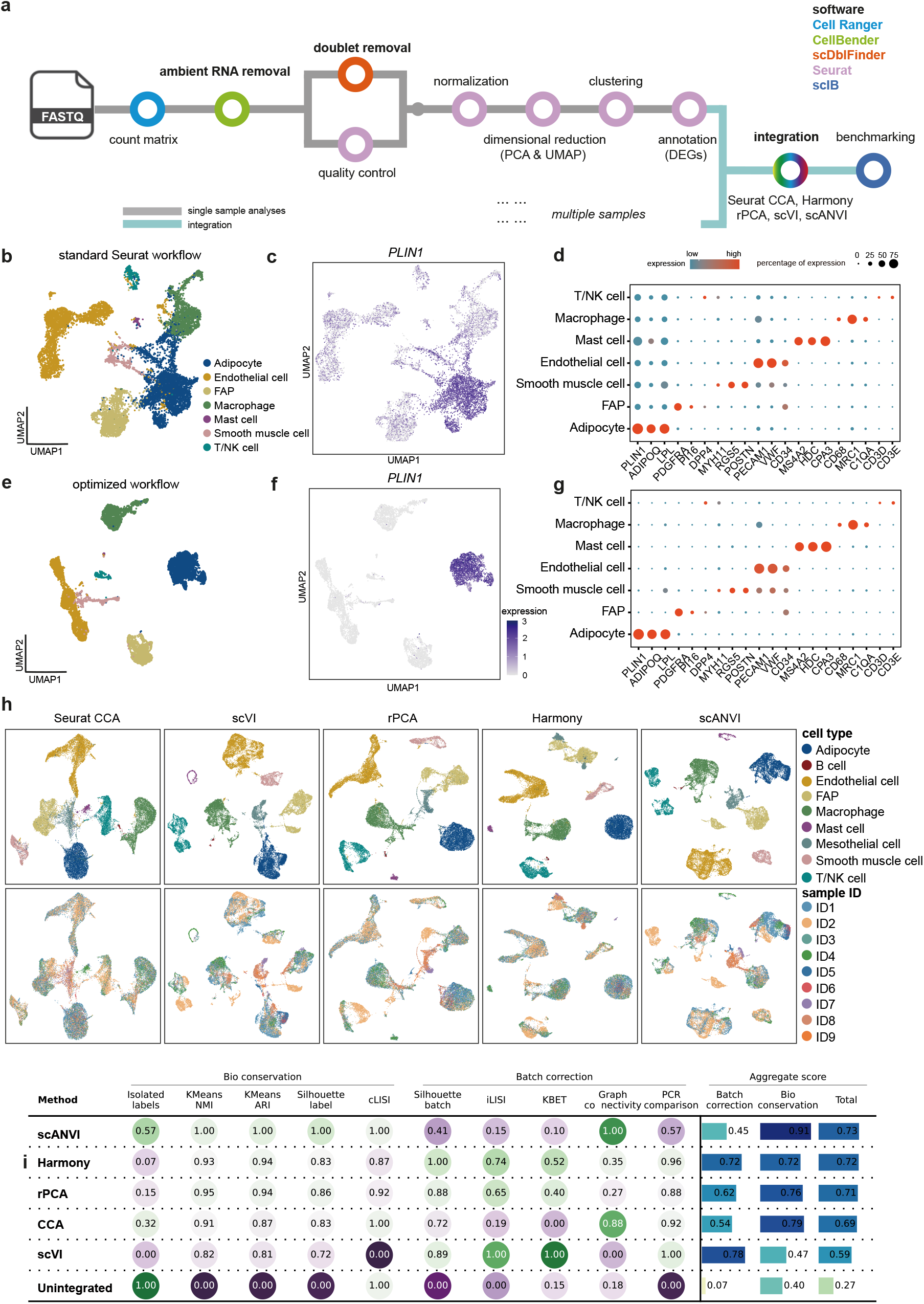
Overview and performance of the optimized snRNA-seq analysis workflow. (a) Schematic representation illustrates the optimized data processing workflow, starting with raw sequencing FASTQ files and proceeding to downstream integration analyses. Key steps include the generation of count matrices using Cell Ranger, removal of ambient RNA contamination with CellBender, doublet identification with scDblFinder, quality control, normalization, dimension reduction (linear by PCA and non-linear by UMAP), annotation of cell types based on differentially expressed genes (DEGs) in Seurat, integration using methods such as Seurat CCA, Harmony, rPCA, scVI, and scANVI, and benchmarking using scIB. (b-d) standard Seurat scRNA-seq pipeline without ambient RNA and doublet removal: (b) UMAP projection of the raw data from one white adipose tissue sample, coloured by annotated cell types: adipocytes, endothelial cells, fibro/adipogenic progenitors (FAP), macrophages, mast cells, smooth muscle cells, and T/NK cells. (c) Expression pattern of the adipocyte marker *PLIN1* in the raw dataset. (d) Dot plot depicts the expression levels of differentially expressed marker genes across cell types. Dot size represents the percentage of cells expressing each marker and colour density reflects the average expression levels. (e-g) after ambient RNA and doublet removal: (e) UMAP projections of cell types coloured by cell type annotations after both ambient RNA and doublet removal. (f) Expression pattern of adipocyte marker *PLIN1* after ambient RNA and doublet removal. (g) Dot plot showing differentially expressed marker genes after both ambient RNA and doublet removal, indicating improved marker specificity and reduced background noise. (h) UMAP representations of the dataset after integration using five batch-effect correction methods: Seurat CCA, scVI, rPCA, Harmony, and scANVI. The first row shows UMAPs coloured by cell type, while the second row shows UMAPs coloured by donor ID. (i) The scIB framework is used for the quantitative assessment of batch-effect correction methods. Metrics are grouped into bio-conservation measures (isolated label performance, KMeans NMI, etc.) and batch correction measures (iLISI, kBET, etc.). Aggregate scores are calculated as a weighted mean (40:60) of batch correction and bio-conservation metrics.

Ambient RNA contamination and doublets commonly occur in the snRNA-seq data during the isolation of the nuclei from individual cells, potentially resulting in incorrect cell type annotation and consequently misleading biological interpretations. We compared the ambient RNA and doublet rates across various samples and tissues and found considerable variability across tissue types (**Supplementary Fig. S1b and c**), with higher ambient RNA rates in liver (up to 90%) and white adipose tissue and higher doublet rates in white adipose tissue and the hypothalamus, highlighting the importance of removing ambient RNA contamination and doublets in snRNA-seq data in these tissues. CellBender was selected for ambient RNA removal after benchmarking against DecontX [14] and SoupX [15] across five datasets (**Supplementary Fig. S1d**). To evaluate the impact of ambient RNA and doublet removal, we compared cell-type annotation results from two independent automatic annoatation methods SingleR [16] and Azimuth [17] before and after applying the optimized pipeline. The results demonstrate a more distinct separation of cell types (**Supplementary Fig. S2a-b**). We further examined the expression of the widely used adipocyte marker gene *PLIN1*, which encodes Perilipin 1, associated with lipid droplet coating, in an illustrated example from white adipose tissue in **Fig. 1b-g**. *PLIN1* was diffusely expressed across nearly all cells prior to the removal of ambient RNA, and in the same time, adipocytes and macrophages were poorly separated (**Fig. 1b-d**). Other adipocytes markers, including *ADIPOQ* and *LPL* demonstrated similar patterns (**Supplementary Fig. S3a**). Additionally, we identified a proportion of doublets expressed both adipocyte marker (*ADIPOQ, LPL*) and macrophage marker (*MRC1*) or endothelial cell marker (*VWF*) (**Supplementary Fig. S3a**).

After the ambient RNA and doublet removal, *PLIN1* expression was exclusively to the adipocyte cluster (**Fig. 1e-g**), reflecting improved cluster separation and enhanced marker genes specificity, as illustrated in UMAP projections, violin plots and density plots (**Supplementary Fig. S3a-d**). We further examined the metabolic pathway enrichment and found that adipocytes exhibited notably higher enrichment for both fatty acid elongation and fatty acid degradation pathways, consistent with their established role in lipid metabolism (**Supplementary Fig. S4**). Dot plot for the differentially expressed marker genes presented refined cell type resolution, including adipocytes (*PLIN1, ADIPOQ, LPL*), fibro/adipogenic progenitors (FAPs) (*PDGFRA, PI16, DPP4*), endothelial cells (*PECAM1, VWF, CD34*), macrophages (*CD68, MRC1, C1QA*), mast cells (*MS4A2, HDC, CPA3*), smooth muscle cells (*MYH11, RGS5, POSTN*), and T cells (*CD3D, CD3E*) (**Fig. 1g**). This improvement also extended to other metabolic tissues we studied (**Supplementary Fig. S5-7**). We observed clearer cell-type clusters of the myofiber (*TTN*) in muscle (**Supplementary Fig. S5)**, the hepatocyte population (*ASGR1*) in liver (**Supplementary Fig. S6)**, and the neuron (*Meg3*) in the hypothalamus (**Supplementary Fig. S7)** post ambient RNA and doublet removal. These steps significantly improve marker specificity and cell-type resolution across metabolic tissues, providing a robust basis for differentially expressed genes that are involved in cell-specific functions and metabolic homeostasis.

A key challenge in multi-sample snRNA-seq studies is the presence of batch effects, which can mask biological signals and introduce technical bias. To tackle this issue, we applied and benchmarked a selection of batch correction/integration methods, including Seurat CCA, Harmony, rPCA, scVI and scANVI across datasets to align cellular profiles across multiple samples and evaluate their performances based on their ability to preserve tissue-specific cell types and minimize batch-associated artifacts. We observed significant sample-specific clustering (**Supplementary Fig. 8a and b**) on UMAP embeddings, indicating technical variability rather than true biological variation in the illustrated adipose tissue dataset before batch correction. After applying various methods, we observed substantial improvements with all integration methods, to varying degrees (**Fig. 1h)**. We adopted the scIB framework which has been benchmarked its performance in evaluation the integration performance, weighted evaluation metrics and found that scANVI was better than other methods in conserving the unique biological transcriptional signatures (score = 0.91) of each cell type across samples with moderate batch correction score (score = 0.45) (**Fig. 1i**), achieving highest aggregated score for integration and clear cell type separation (**Fig. 1h**). Harmony ranked the second-best integration method in this dataset with both relative high score in batch correction (score = 0.72) and biological signal conservation (score = 0.72) (**Fig. 1i**). Moreover, we found Harmony consistently performed well for the liver (**Supplementary Fig. S9**), human muscle and the hypothalamus [18]. Taken together, our benchmarking of integration methods highlights the need for careful method selection for processing multi-sample snRNA-seq data and most appropriate method may vary depending on the specific data, and Harmony has emerged as an effective tool across tissues, making it the first choice for dataset integration.

Our study aims to delineate a standardized upstream analytical workflow for snRNA-seq analysis in metabolically active tissues, including white adipose tissue, muscle, liver and the hypothalamus, to achieve accurate cell-type clustering, annotation and functionally relevant cellular insights by addressing technical challenges including ambient RNA and doublet contamination and rigorously benchmarked the integration of data from multiple samples. Our workflow enhances the resolution of cell clusters and ensures that subsequent analyses are based on high-quality, biologically meaningful data, facilitating exploration of tissue-specific mechanisms underlying metabolic homeostasis and diseases.

We found substantial tissue-specific variability in ambient RNA and doublet rates. Liver and adipose tissue showed notably higher ambient RNA rates compared to other tissues. These variations likely arise from differences in tissue architecture, cell size, and nuclear RNA content, significantly impacting data quality and analytical approaches. Understanding these tissue-specific characteristics is crucial for optimizing preprocessing parameters and ensuring accurate biological interpretation. A major advancement in our work is the integration of established tools, specifically optimized for metabolic tissues in addressing ambient RNA contamination and doublet removal. With thousands of packages available in the single-cell field, selecting appropriate tools can be challenging for biologists. Our workflow addresses this challenge by carefully benchmarking these tools and showed CellBender and scDblFinder outperform other packages in metabolic tissue analyses. Ambient RNA contamination, originating from extracellular RNA that enters nuclei suspensions, can result in non-specific gene expression across cell types, hence obscuring true biological signals. We showed an ambient RNA removal step can effectively minimize this noise, allowing accurate identification of cell-type specific markers. *PLIN1*, a hallmark adipocyte marker gene, was shown to be as exclusively expressed in adipocytes only after ambient RNA removal, as well as other marker gene expression in muscle, liver and the hypothalamus, confirming the necessity of this preprocessing step for accurate cell type annotation. This improved cell-type resolution (eg. adipocytes) and identification of biologically relevant pathways is crucial for understanding metabolic homeostasis and disease states such as differences in the white adipose tissue in obesity and type 2 diabetes, where precise characterization of cellular subpopulations is essential for unravelling tissue remodeling and inflammation processes.

The integration of multi-sample datasets presented a key technical challenge that we addressed by benchmarking several batch correction methods, including Seurat CCA, Harmony, rPCA, scVI, and scANVI. Effective batch correction is crucial in multi-sample studies to reduce technical variability without compromising biologically meaningful cell-type differences, representing the most challenging analytical step in large consortium/atlas [3, 10]. To address the difficulty of selecting the most appropriate integration method, we adopted the scIB framework, which has been benchmarked for evaluating integration performance. Its weighted evaluation metrics provide a systematic approach to method selection. We found Harmony demonstrate strong performance in our presented data whereas scVI also showed commendable results in previously published studies on the hypothalamus [19] and white adipose tissue [6]. Notably, users can adjust the parameters in the batch correction methods to enhance the compatibility for their datasets followed by scIB. For example, users can set varying *theta* values for different covariates in the Harmony method. Our benchmarking analysis emphasizes the necessity of choosing suitable integration methods customized to the complexity and composition of each tissue, reinforcing the importance of a standardized workflow to maintain biological integrity across samples.

While our optimized workflow significantly enhances snRNA-seq data quality and analytical reliability, there are several limitations. First, the workflow requires basic computational resources and bioinformatics knowledge, meaning researchers should have a fundamental understanding of R or Python programming. To help with this, we have developed an open-access tutorial website and recommend cloud-based computing options to facilitate broader adoption. Second, tissue-specific variability in nuclear RNA content and cellular composition necessitates customized preprocessing parametres, as demonstrated by the distinct ambient RNA and doublet rates observed across different metabolic tissues. Although scIB provides an integrated evaluation framework, careful examination is still required to avoid overcorrection or loss of biological variability. Third, this workflow has been primarily optimized for snRNA-seq data, and its adaptability to other omics datasets, such as single-cell ATAC-seq, remains to be explored. Future improvements, including multi-omics integration and AI-driven analysis tools will be essential for expanding its applications across diverse research settings. Last but not the least, we acknowledge that we have not integrated downstream analyses into the workflow, given the complexity and diversity of biological questions that need to be addressed.

In summary, our study underlines the critical role of a standardized workflow in enhancing data quality and analytical reliability in snRNA-seq of metabolic tissue by combining rigorous preprocessing, cell-type clustering and annotation, and robust integration methods. By providing an open-access tutorial website (https://metabomicslab.github.io/snRNAseq-analysis-workflow/) and snakemake pipeline, we aim to make these analytical tools more accessible to the broader research community. This will broaden single nuclei transcriptomics application in metabolic biology to elucidate cellular heterogeneities and tissue-specific mechanisms and forming a strong foundation for future precision medicine targeting cell-specific functions.

## Data availability

All raw sequencing data are publicly available in GSE217677, GSE225700, GSE202379, PRJNA772047 and PRJNA771932 and all the re-analysed Cell Ranger output were available on Zenodo: https://zenodo.org/records/14725531.

## Code availability

All scripts used in the current study are available on GitHub: https://metabomicslab.github.io/snRNAseq-analysis-workflow/ and an integrated snakemake pipeline for the upstream analyses (https://zenodo.org/records/14725531)

## Acknowledgements

This study was funded by the National Key Research and Development Program of China, National Natural Science Foundation of China and a new PI Start up grant of Fudan University to GW. The computations in this research were performed using the CFFF platform of Fudan University and Medical Science Data Center of Fudan University. The schematic workflow in **Supplementary Fig. S1a** were adapted and created with BioRender.com (Agreement number:PV27N9UNFF).

## Author Contributions

Conceptualization, GW; Methodology, GW and PD; Investigation, GW and PD; Data analysis, PD and SD; Writing, PD and GW; Supervision, GW

